# Long-read sequencing across the *C9orf72* ‘GGGGCC’ repeat expansion: implications for clinical use and genetic discovery efforts in human disease

**DOI:** 10.1101/176651

**Authors:** Mark T. W. Ebbert, Stefan Farrugia, Jonathon Sens, Karen Jansen-West, Tania F. Gendron, Mercedes Prudencio, lan J. McLaughlin, Brett Bowman, Matthew Seetin, Mariely DeJesus-Hernandez, Jazmyne Jackson, Patricia H Brown, Dennis W. Dickson, Marka van Blitterswijk, Rosa Rademakers, Leonard Petrucelli, John D. Fryer

## Abstract

**Background:** Many neurodegenerative diseases are caused by nucleotide repeat expansions, but most expansions, like the *C9orf72* ‘GGGGCC’ (G_4_C_2_) repeat that causes approximately 5-7% of all amyotrophic lateral sclerosis (ALS) and frontotemporal dementia (FTD) cases, are too long to sequence using short-read sequencing technologies. It is unclear whether long-read sequencing technologies can traverse these long, challenging repeat expansions. Here, we demonstrate that two long-read sequencing technologies, Pacific Biosciences’ (PacBio) and Oxford Nanopore Technologies’ (ONT), can sequence through disease-causing repeats cloned into plasmids, including the FTD/ALS-causing G_4_C_2_ repeat expansion. We also report the first long-read sequencing data characterizing the *C9orf72* G_4_C_2_ repeat expansion at the nucleotide level in two symptomatic expansion carriers using PacBio whole-genome sequencing and a no-amplification (No-Amp) targeted approach based on CRISPR/Cas9.

**Results:** Both the PacBio and ONT platforms successfully sequenced through the repeat expansions in plasmids. Throughput on the MinION was a challenge for whole-genome sequencing; we were unable to attain reads covering the human *C9orf72* repeat expansion using 15 flow cells. We obtained 8x coverage across the *C9orf72* locus using the PacBio Sequel, accurately reporting the unexpanded allele at eight repeats, and reading through the entire expansion with 1324 repeats (7941 nucleotides). Using the No-Amp targeted approach, we attained >800x coverage and were able to identify the unexpanded allele, closely estimate expansion size, and assess nucleotide content in a single experiment. We estimate the individual’s repeat region was >99% G_4_C_2_ content, though we cannot rule out small interruptions.

**Conclusions:** Our findings indicate that long-read sequencing is well suited to characterizing known repeat expansions, and for discovering new disease-causing, disease-modifying, or risk-modifying repeat expansions that have gone undetected with conventional short-read sequencing. The PacBio No-Amp targeted approach may have future potential in clinical and genetic counseling environments. Larger and deeper long-read sequencing studies in *C9orf72* expansion carriers will be important to determine heterogeneity and whether the repeats are interrupted by non-G_4_C_2_ content, potentially mitigating or modifying disease course or age of onset, as interruptions are known to do in other repeat-expansion disorders. These results have broad implications across all diseases where the genetic etiology remains unclear.

## Background

Many neurodegenerative diseases, including Huntington’s disease [1–4], spinocerebellar ataxias [1,2], frontotemporal dementia (FTD) [3], and amyotrophic lateral sclerosis (ALS) [3] can be caused by nucleotide repeat expansions [1] that are historically challenging to sequence [4,5]. Repeat expansions are a specific multi-nucleotide DNA sequence that is repeated (i.e., expanded) significantly more times than normal. In 2011, a *C9orf72* ‘GGGGCC’ (G_4_C_2_) repeat expansion was discovered [3,6] that causes approximately 34% and 26% of familial ALS and FTD cases, respectively [7]. This finding genetically linked ALS and FTD, generating an exciting opportunity to better understand the etiology of both diseases, and potentially develop a therapeutic approach. Individuals with ALS and FTD caused by the G_4_C_2_ expansion generally have hundreds to thousands of G_4_C_2_ repeats [8], while healthy individuals typically have between 2 and 30 G_4_C_2_ repeats [6,9], though a precise cutoff for pathogenicity is unclear [9]. Additional diseases caused by repeat expansions include Fuch’s disease [10], myotonic dystrophy [11], Friedreich’s ataxia [12], and Fragile X syndrome [13], among others, demonstrating the breadth of diseases caused by such expansions. Revealing the underlying etiology of these diseases, and discovering additional repeat expansions that directly cause or modify disease, or modify risk for disease, will likely be accelerated through long-read sequencing technologies capable of characterizing at least major portions of the repeat; characterizing these repeats at the nucleotide level will help determine, for example, whether the repeat is interrupted and whether such interruptions mitigate disease, as in other neurodegenerative disorders [14–17].

It is unclear whether third-generation long-read sequencing platforms such as Pacific Biosciences’ (PacBio; RS II and Sequel) and Oxford Nanopore Technologies’ (ONT; MinION) can traverse these challenging disease-causing repeats, nor is there a report of nucleotide-level sequencing data in a *C9orf72* repeat expansion carrier. Likewise, it is unclear whether the *C9orf72* repeat expansion is pure G_4_C_2_ repeat in affected carriers, or whether it is interrupted by non-G_4_C_2_ sequence. The *C9orf72* G_4_C_2_ expansion may be the most challenging repeat to sequence, given its extreme length, “pure” GC content [4,5], and propensity to form G-quadruplexes in both RNA [18,19] and DNA [19].

Here, we demonstrate that both PacBio and ONT sequencing platforms can sequence through repeats cloned into plasmids, including the spinocerebellar ataxia type 36 (SCA36) disease-causing ‘ GGCCTG’ repeat expansion [20] and the FTD- and ALS-causing G_4_C_2_ repeat expansion. We further report long-read sequencing data from the *C9orf72* G_4_C_2_ repeat expansion at the nucleotide level in two symptomatic expansion carriers using both whole-genome and no-amplification (No-Amp) targeted sequencing [21,22] on the PacBio Sequel. Our findings indicate that long-read sequencing is well suited to characterizing repeat expansions and that this technology has potential to accelerate future genetic discovery efforts across a broad range of diseases that may involve repeat expansions. These technologies may also have potential in clinical and genetic counseling environments for repeat-expansion and other structural variant disorders, generally. Structural mutations, and repeat expansions specifically, are challenging for short-read technologies. Thus, long-read sequencing technologies may be ideal for discovering new disease-causing or disease-modifying repeat expansions that have escaped detection with conventional short-read sequencing.

## Results

### PacBio RS II and ONT MinlON sequence through repeats cloned into plasmids

To generally assess the PacBio and ONT sequencing platforms, we cloned the SCA36 ‘GGCCTG’ (Figure 1b) and *C9orf72* G_4_C_2_ (Figures 1c and 1d) repeat expansions into plasmids and sequenced them on the PacBio RS II and ONT MinION (Figure 2). We also included an EGFP-containing plasmid without a repeat expansion, as a control (Figure 1a). The SCA36 plasmid was included as a secondary control because it has 1/6^th^ lower GC content than the *C9orf72* G_4_C_2_ repeat; we anticipated that the G_4_C_2_ repeat may be too challenging for these technologies, as sequencing GC-rich regions has historically been challenging for any technology [4,5]. We aimed to construct a plasmid containing 62 ‘GGCCTG’ repeats, and two plasmids containing approximately 423 (C9-423) and 774 (C9-774) G_4_C_2_ repeats, respectively. Because these long repeat sequences are unstable, most bacterial colonies contained fewer than the targeted number of repeats (Supplemental figure 1).

**Figure 1.**
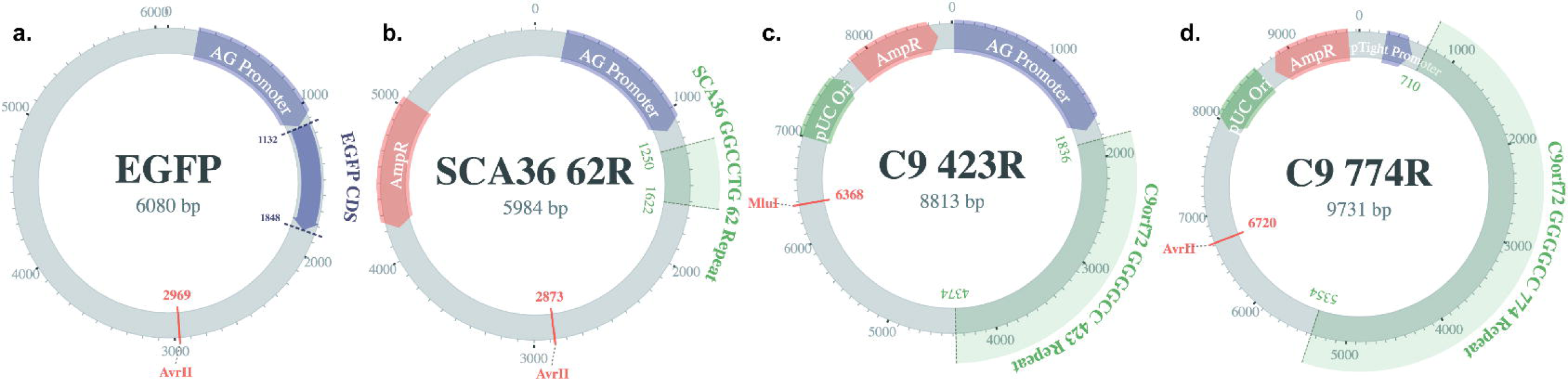
Schematic diagrams for plasmids used to test PacBio and ONT long-read sequencing technologies. To minimize biases when comparing the PacBio RS II and ONT MinION, we constructed four plasmids, including three repeat-containing plasmids and a non-repeat-containing plasmid. Each plasmid map identifies estimated plasmid size, and the location and size of the repeat within the plasmid, **(a)** The first plasmid did not contain a repeat, as a control, but instead included the *EGFP* gene. The *EGFP* plasmid was linearized at position 2969 with the Avril restriction enzyme, **(b)** We also constructed a plasmid with 62 repeats of the spinocerebellar ataxia type 36 (SCA36) ‘GGCCTG’ repeat, which was linearized at position 2873 with Avril, **(c)** A third plasmid contained 423 *C9orf72* ‘GGGGCC’ repeats, and was linearized at position 6368 with Mlul to maximize non-repeat sequence both up and downstream of the plasmid, thus avoiding bias against reads in either direction; allowing the repeat to be too close to either end could compromise sequencing or downstream analyses, **(d)** We included an additional plasmid with 774 *C9orf72* ‘GGGGCC’ repeats to simulate the expansion size found in ALS- or FTD-affected expansion carriers. While 774 repeats is dramatically smaller than the expansion found in many affected carriers, it was the largest we were able to construct reliably, because these repeats are unstable in bacteria. Additionally, while we targeted the number of specified repeats for each plasmid, most colonies contained fewer than the targeted repeats because repeats are generally unstable in bacteria (Supplementary Figure 1). Thus, the targeted number of repeats serves as an estimated maximum number of repeats.

**Figure 2.**
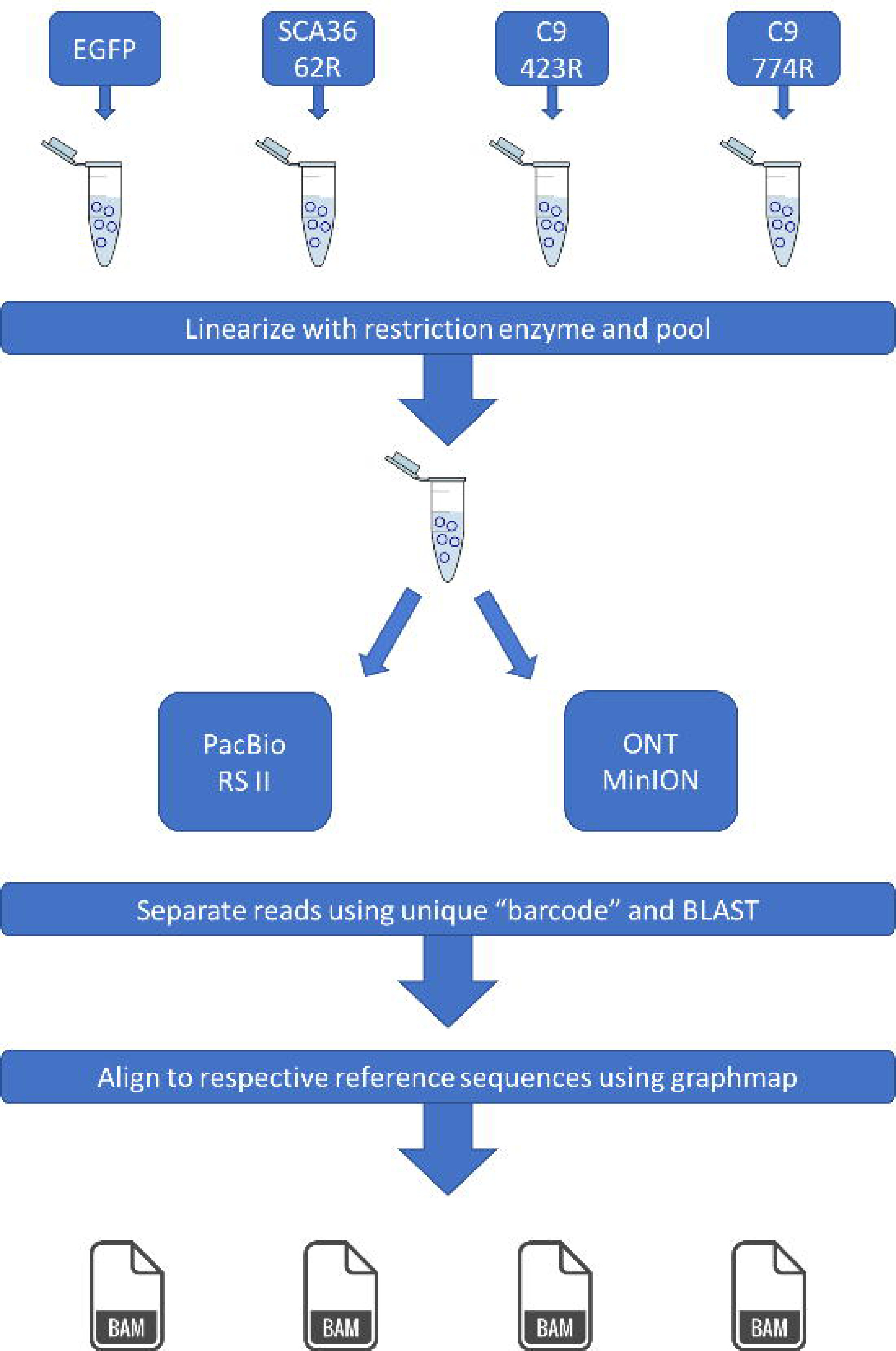
Workflow for linearizing, pooling, and sequencing plasmids on the PacBio RS II and ONT MinION long-read platforms. Each plasmid was cut with the restriction enzyme identified in the respective plasmid maps (Figure 1), and at the specified location. After linearizing each plasmid independently, the plasmids were pooled and cleaned. We then sequenced the same pool on the PacBio RS II and Oxford Nanopore Technologies’ (ONT) MinION using their respective library preparation protocols. After sequencing, reads from each plasmid were identified using BLAST and then aligned to their respective reference sequences using graphmap, as preparation for downstream comparisons.

Of the reads that aligned to the respective plasmid sequences, there were 46213, 67339, 9012, and 11535 PacBio RS II subreads for EGFP, SCA36, C9-423, and C9-774, respectively (Figure 3). Similarly, there were 26736, 39059, 8276, and 8720 ONT MinION reads for the same plasmids, respectively. Read length distributions for the ONT MinION were much tighter than those from the PacBio RS II, particularly for the C9-423 and C9-774 plasmids. In general, both platforms had a single mode (i.e., peak) and had similar median read lengths (Figure 3), with the exception of the PacBio RS II C9-774 read length distribution. PacBio RS II median read lengths for EGFP, SCA36, C9-423, and C9-774 were 5854, 5676, 6396, and 5714, respectively, while ONT MinION median read lengths were 5879, 5660, 7024, and 7313, respectively (Figure 3). Expected maximum read lengths for each plasmid were 6080 (EGFP), 5984 (SCA36), 8813 (C9-423), and 9731 (C9-774) (Figure. 1, 3). Approximately 30.2% (EGFP), 19.2% (SCA36), 6.9% (C9-423), and 3.2% (C9-774) PacBio RS II reads were longer than the expected maximum length, whereas only approximately 0.5%, 0.3%, 0.3%, and 0.1% ONT MinION reads were longer than the expected maximum length.

**Figure 3.**
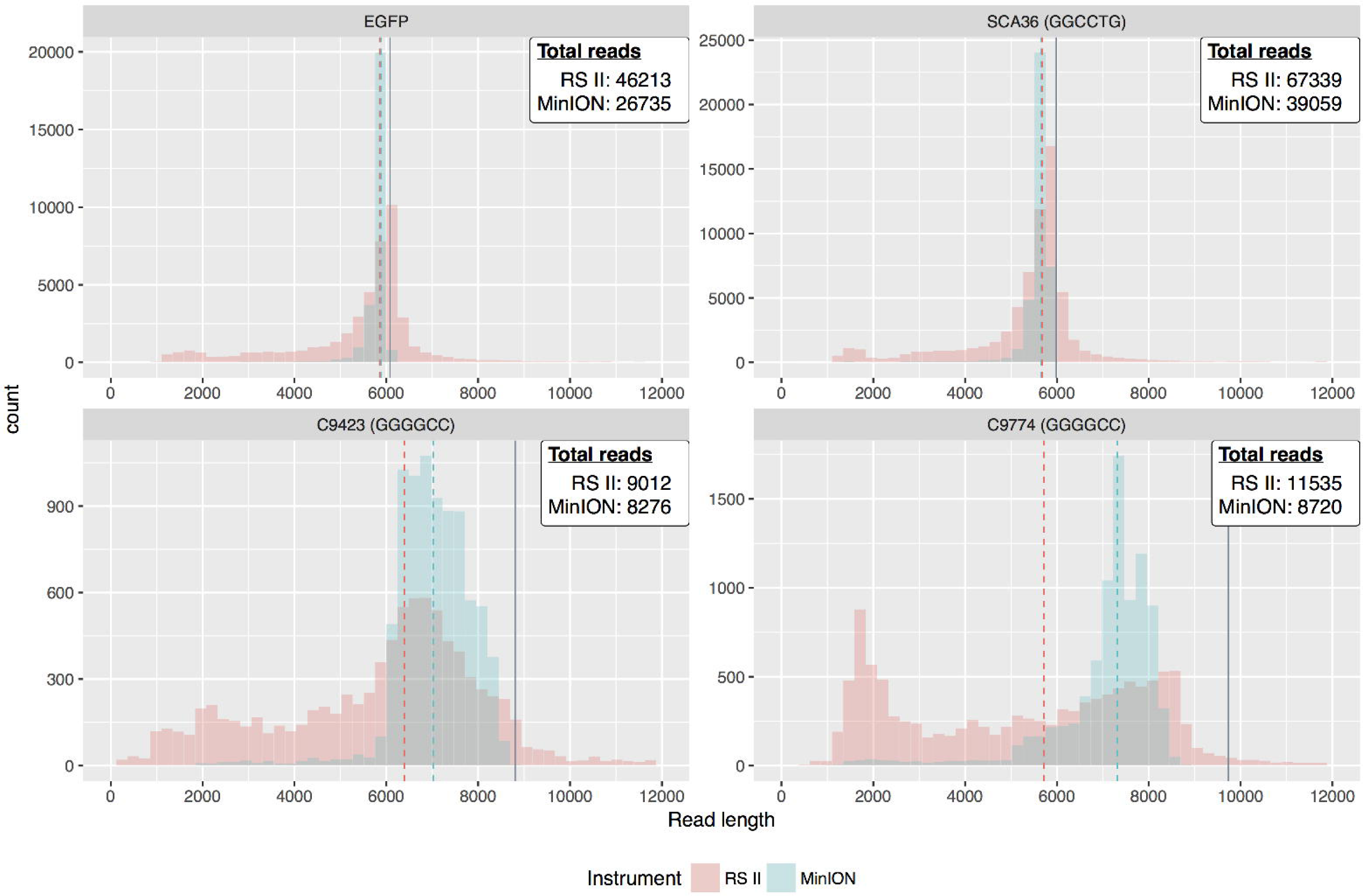
Both the PacBio RS II and ONT MinlON successfully sequence through repeats, but the RS II had more variable read lengths. After selecting only those reads that could be clearly identified for each plasmid (described in Figure 1), there were 46213, 67339, 9012, and 11535 PacBio RS II reads for *EGFP*, SCA36, C9-423, and C9-774, respectively. Likewise, there were 26735, 39059, 8276, and 8720 ONT MinlON reads for the same respective plasmids. The PacBio RS II generally had more reads, but read length distributions are much tighter for the ONT MinlON across all four plasmids, and more closely resemble expected read lengths. The median read length for each instrument is indicated by dashed lines, and the expected maximum read length is indicated by a solid gray line. Expected maximum read lengths for each plasmid were 6080 *(EGFP)*, 5984 (SCA36), 8813 (C9-423), and 9731 (C9-774). Because these long repeat sequences are unstable in plasmids, however, most bacterial colonies contained fewer than the targeted number of repeats (Supplemental figure 1). Thus we expect the read sizes to vary. The additional PacBio RS II read variability may be related to library preparation.

We also compared how well the platforms sequenced specifically through the repeat regions, where we first compared repeat length distributions. Both platforms produced highly similar distributions for all plasmids (Figure 4), but the repeat lengths varied widely within each plasmid, as expected based on gel intensity curves (Supplemental figure 1). The C9-423 repeat length distribution is more variable than even the C9-774, perhaps because the C9-774 plasmid backbone is more tolerant of the repeat. The median repeat lengths are highly similar for the PacBio RS II and ONT MinION (Figure 4), where the median repeat lengths (measured by repeat number, not bases) for the PacBio RS II were 35,148, and 395 for SCA36, C9-423, and C9-774, respectively, while median repeat lengths for the ONT MinlON were 37,172, and 406, respectively. Repeat lengths for the SCA36 plasmid were confirmed by Sanger sequencing, estimating approximately 37 repeats, before sequence traces became indeterminate (Supplemental figure 2). We also assessed the percentage of reads that extended through the repeats to assess whether the innate characteristics of the repeat affected sequencing performance. Approximately 95.9%, 66.8%, and 43.8% of PacBio RS II reads successfully extended through the repeat for SCA36, C9-423, and C9-774, respectively, while 99.5%, 97.7%, and 83.5% of ONT MinION reads extended through, respectively.

**Figure 4.**
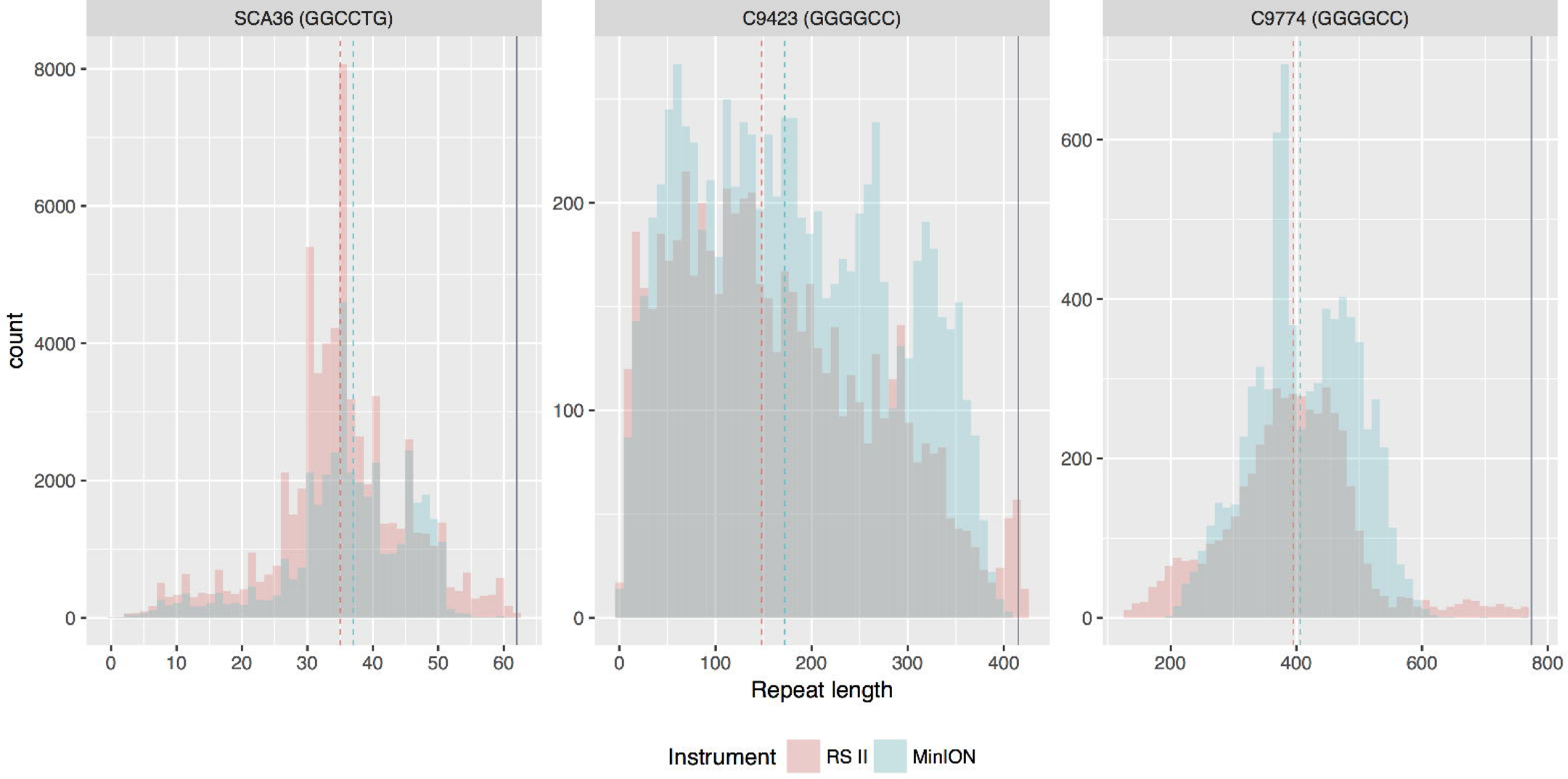
Repeat length distributions for the PacBio RS II and ONT MinlON were highly concordant. Both platforms produced highly similar distributions for all plasmids, but the repeat lengths varied widely within each plasmid, as expected based on gel intensity curves (Supplemental figure 1). The C9-423 repeat length distribution is more variable than even the C9-774, perhaps because the C9-774 plasmid backbone is more tolerant of the repeat. The median number of repeats for the PacBio RS II were 35,148, and 395 for SCA36, C9-423, and C9-774, respectively, while median repeat lengths for the ONT MinlON were 37,172, and 406, respectively. The percentage of reads that extended through the SCA36, C9-423, and C9-774, repeats were approximately 95.9%, 66.8%, and 43.8% for the PacBio RS II, respectively, while 99.5%, 97.7%, and 83.5% of ONT MinlON reads extended through, respectively.

Base calling accuracy through the C9-774 repeat region was dramatically different between the PacBio RS II and ONT MinION for the *C9orf72* repeat. After aligning the repeat region for individual reads to the expected repeat sequence of the same length, using the global Needleman-Wunsch algorithm [23], the median PacBio RS II error rate for C9-774 was 7.4%, while the median ONT MinION error rate for C9-774 was 47.3%. The PacBio RS II consensus sequence (Supplemental data 1) contained 774 repeats in the C9-774 plasmid, attaining approximately 99.8% accuracy. The ONT MinION consensus sequence (Supplemental data 2) also contained 774 repeats in the C9-774 plasmid, but was only approximately 26.6% accurate, because many guanines and cytosines were erroneously called as adenine. Thus, in the ONT MinION consensus sequence, guanines and cytosines were represented as mixed nucleotides in the consensus sequence (e.g., R representing G or A, and M representing C or A). Exactly 553 (71.4%) of the 774 ONT MinION repeats were represented as either RRRRCM orRRRRMC.

### PacBio Sequel successfully sequences through the *C9orf72* repeat expansion in affected carriers

Sequencing repeat expansions cloned into plasmids demonstrated that both platforms are capable of sequencing through these challenging repeats, but demonstrating on a human expansion carrier is essential to determining whether long-read technologies can characterize the repeats in their entirety at the nucleotide level, and also to determine whether the technologies are suitable for discovering new disease-causing or disease-modifying repeat expansions. We identified and confirmed two *C9orf72* G_4_C_2_ repeat expansion carriers through fluorescent PCR fragment analysis, repeat-primed PCR, and Southern blotting [24] using cerebellar tissue. The fluorescent PCR analysis demonstrated the individuals carried two and eight repeats in the unexpanded allele, respectively (Figure. 5 a, d). We then determined the individuals were expansion carriers through repeat-primed PCR (Figure. 5 b, e), and estimated the expansion sizes by Southern blot (Figure. 5 c, f). The most abundant expansion sizes, averaged across multiple Southern blots, indicate repeat sizes of approximately 1083 repeats (8.8kb, including flanking sequence) and 1933 repeats (13.9kb), respectively.

**Figure 5.**
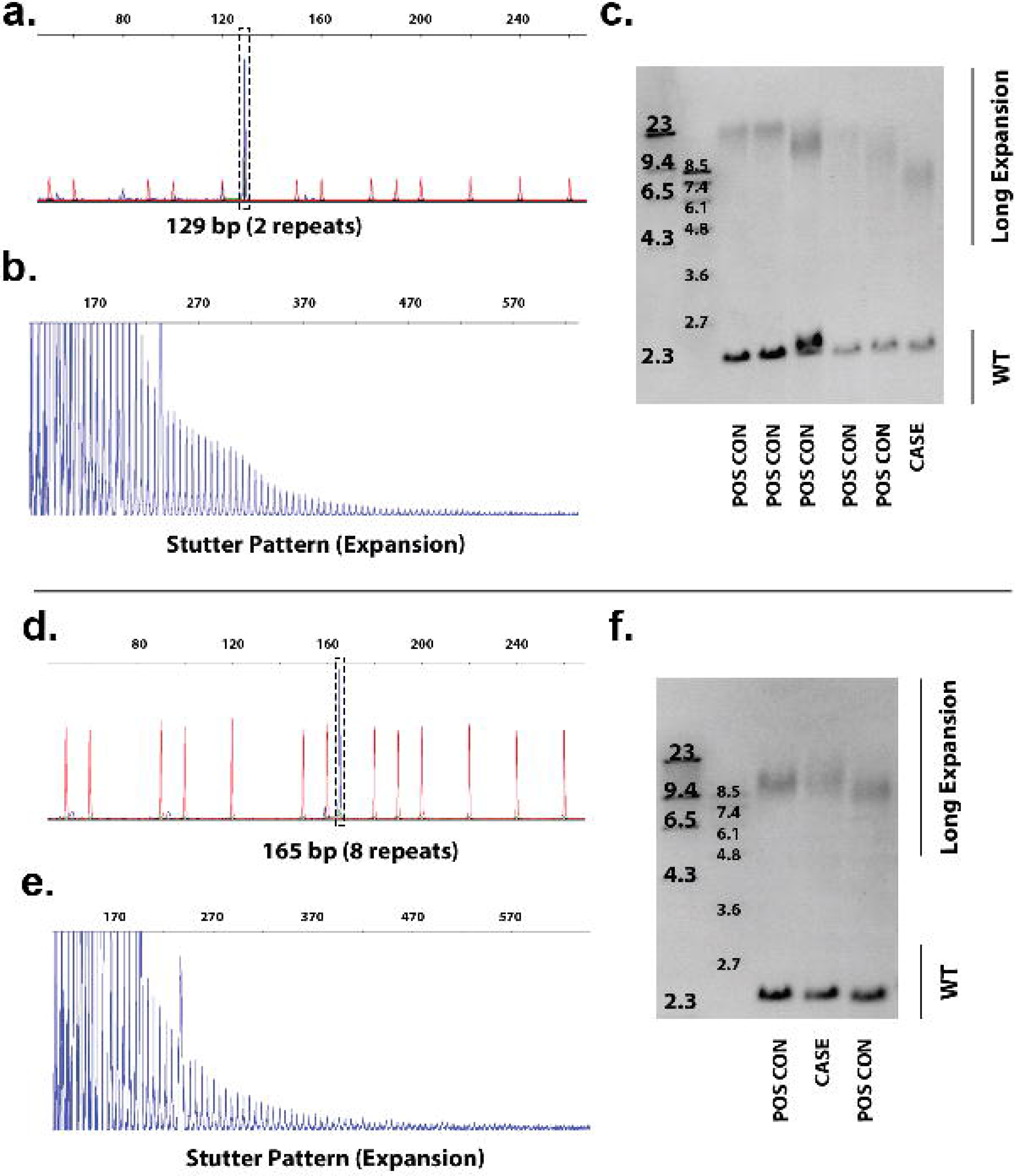
Characterization of the affected *C9orf72* repeat expansion carriers using standard methodologies. **(a, d)** We first performed fluorescent PCR to determine the individual’s non-pathogenic repeat size. Genomic DNA was PCR-amplified with genotyping primers and one fluorescently labeled primer. Fragment length analysis of the PCR product was then performed on an ABI3730 DNA analyzer and visualized using GeneMapper software. A peak is observable at 129 bp (a) and 165 bp (d), indicating that the non-pathogenic alleles for samples 1 and 2 contain two and eight repeats, respectively. A single peak also indicates that the individual is either homozygous for the given allele, or also has an expansion, **(b, e)** To determine whether the individuals had a repeat expansion, we performed a repeat-primed PCR analysis. PCR products of a repeat-primed PCR were separated on an ABI3730 DNA analyzer and visualized by GeneMapper software, showing a stutter amplification characteristic for a *C9orf72* repeat expansion. This does not indicate expansion size, however, **(c, e)** After determining the individuals were expansion carriers, we performed a Southern blot to estimate the size. The Southern blots reveal a long repeat expansion in other individuals for whom cerebellar tissue was available, including positive controls (POS CON; lanes 1-5, and 1 and 3, respectively) and our patients of interest (CASE; lanes six and two, respectively). DIG-labeled DNA Molecular Weight Markers (Roche) are shown to estimate the repeat expansion’s size. Measurements were based on multiple separate Southern blots for each case; for simplicity one representative Southern blot is shown. The most abundant expansion size in samples 1 and 2 are estimated around 1083 (8.8kb) and 1933 repeats (13.9kb), respectively. The smears ranged widely, demonstrating the heterogeneity (i.e., mosaicism) of this repeat expansion within a small piece of tissue. This demonstrates the importance of additional long-read sequencing studies to characterize the repeat at the nucleotide level.

We then performed whole-genome sequencing on the sample with the longer repeat (sample 2) on both the PacBio Sequel and the ONT MinION. We transitioned to the PacBio Sequel from the RS II because the Sequel’s higher throughput is more amenable to whole-genome sequencing for large genomes. We purified high molecular-weight cerebellar DNA for sample 2 and generated approximately 7x median genome-wide coverage, and 8x coverage across the *C9orf72* repeat locus from five PacBio Sequel SMRT cells. We sequenced the same sample on the ONT MinION, generating approximately 3x median coverage, and 2x across the *C9orf72* repeat locus from 15 flow cells. All ONT MinION flow cells passed quality control before loading the library, with >1000 active pores.

Of the two ONT MinION reads, neither covered an expanded allele, and the repeat region for only one of the reads could be clearly defined. We excluded the read for which we could not clearly define the repeat region. Where the human reference genome (hg38) contains three G_4_C_2_ repeats (Figure 6a), the read for which we could clearly define the repeat region had a total of 41 nucleotides within the repeat region (gain of 25 nucleotides compared to hg38); this equates to approximately seven total repeats (Supplemental figure 3b; Supplemental data 3). While hg38 contains three repeats (Supplemental figure 3a), this does not accurately represent what is observed in the population, as the most common non-pathogenic allele is two repeats followed by eight repeats [3]. An allele with three repeats was not observed in the population [3]. The ONT MinION’s measurement of seven repeats closely resembles the eight repeats measured by our fluorescent PCR fragment analysis (Figure 5d). The ONT MinION was only able to sequence the non-mutant allele with 15 flow cells in this study.

**Figure 6.**
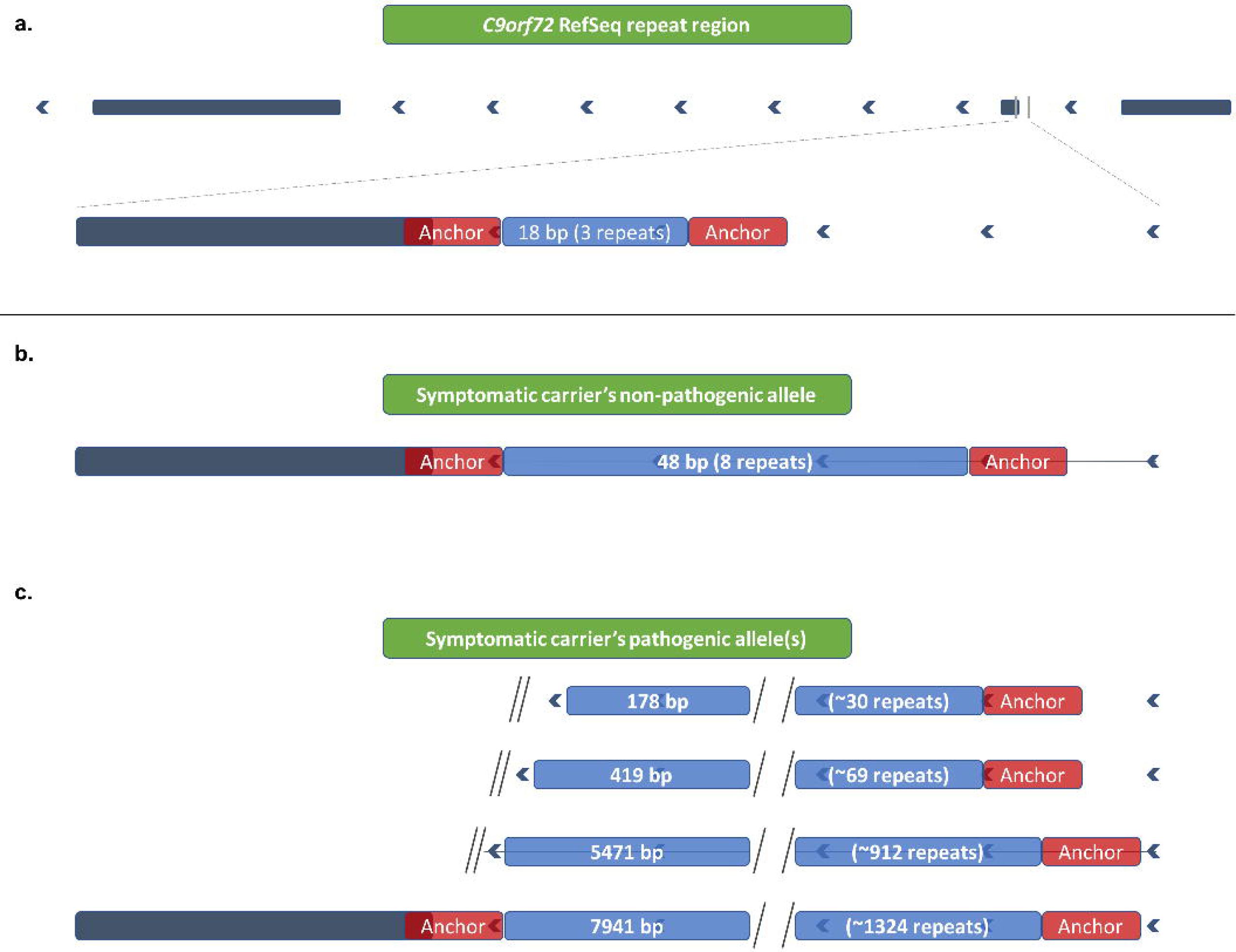
PacBio Sequel reads traverse the repeat region for pathogenic and non-pathogenic alleles. The PacBio Sequel sequenced through both pathogenic and non-pathogenic alleles, demonstrating the platform is capable of characterizing repeat expansions. All of these reads were first aligned by graphmap, and then hand curated to determine the repeat region, **(a)** The human genome reference sequence (hg38) contains three G_4_C_2_ repeats (18 nucleotides). We identified specific “landmarks” before and after the repeat region in the reference sequence to properly locate the repeat region in the reads, and to hand curate the alignments. Landmarks are identified by red bars adjacent to the repeat region, **(b)** We obtained four PacBio Sequel reads covering the eight-repeat sequence, spanning 48 nucleotides. There was a net gain of 29 nucleotides within the defined repeat region, which equates to approximately 5 additional repeats; this concurs with our fragment analysis (Figure 5a). **(c)** We also obtained four reads that covered an expanded allele, one of which bridged the entire repeat expansion, with approximately 1324 repeats (7941 nucleotides). The other three reads ended before bridging the repeat region, where one captured approximately 30 repeats (178 nucleotides), another captured approximately 69 repeats (419 nucleotides), and the third captured approximately 912 repeats (5471 nucleotides).

**Figure 7.**
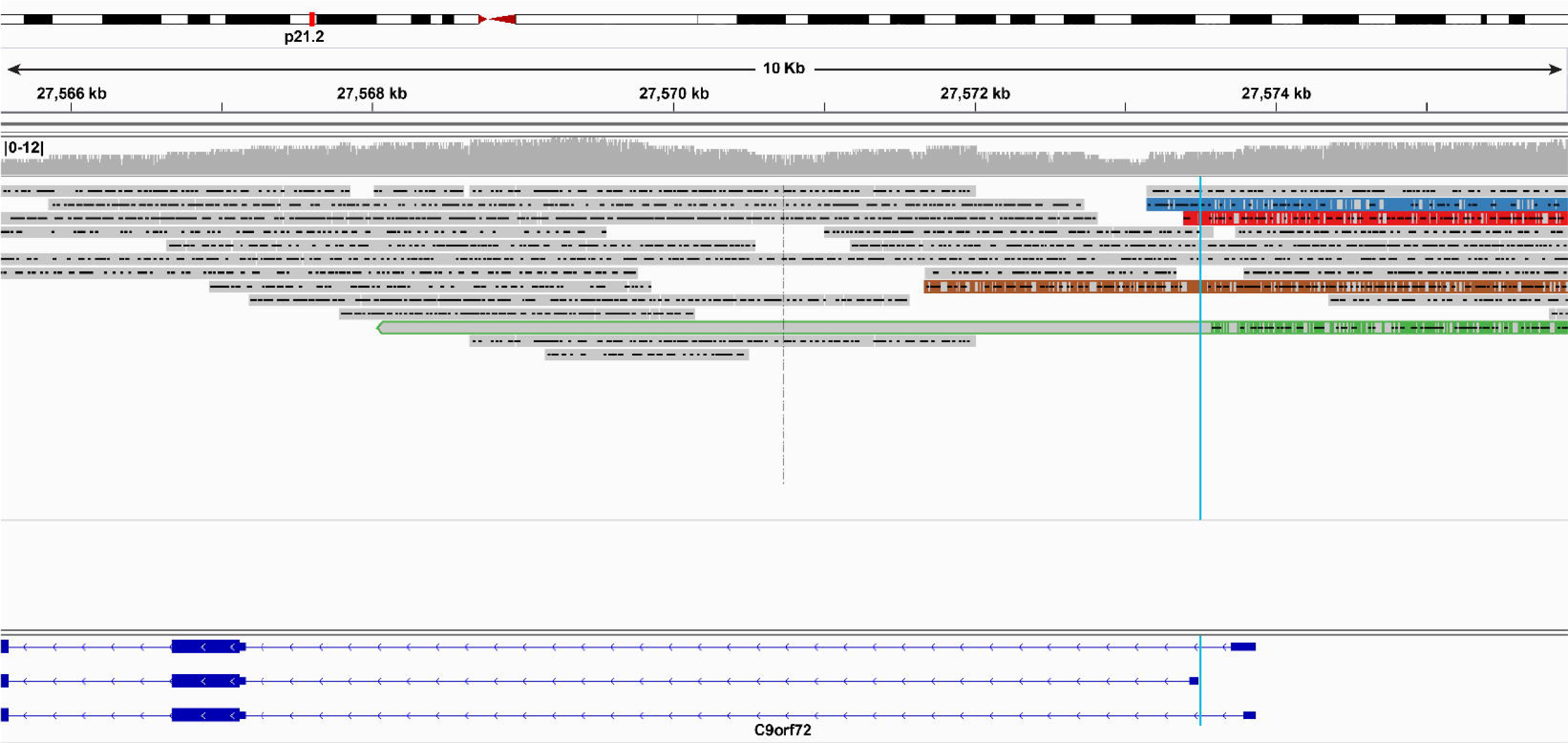
Whole-genome PacBio Sequel reads aligned to hg38. Whole-genome reads generated using the PacBio Sequel were aligned to human reference genome hg38 using graphmap. We attained 7x genome-wide median coverage and 8x across the *C9orf72* repeat locus. Four reads were from the individual’s wild-type allele of eight repeats, while the other four, were expanded. Three of the four reads capturing an expanded allele did not bridge the entire repeat region, where one captured 178 nucleotides in the repeat region (approximately 30 repeats; red), another captured 419 nucleotides (approximately 69 repeats; blue), and the third captured 5471 nucleotides (approximately 912 repeats; green). The read capturing 419 nucleotides may have bridged the repeat because the end of the read closely matches the sequence adjacent to the repeat region, but was ambiguous (Supplemental figure 3d). The final read spanned the entire repeat with 7941 nucleotides (approximately 1324 repeats; brown), which falls easily within the Southern blot’s range (Figure 5f). Soft-clipped nucleotides—nucleotides at the end of a read that did not align to the reference—are shown for all reads, and are outlined in green for the read capturing 912 repeats. The approximate location for the repeat expansion is marked by the light-blue lines. A histogram showing read depth per nucleotide is included near the top of the figure. Alignments were visualized using the Integrative Genomics Viewer (IGV).

### Whole-genome PacBio Sequel sequencing identifies repeat expansion and characterizes repeat length

To date, researchers have relied on Southern blots to measure an individual’s repeat expansion size, but technologies have limited our ability to characterize nucleotide content, which may have critical implications on disease etiology, age of onset, duration, and other clinical phenotypes. The whole-genome PacBio Sequel reads enabled us to generally characterize repeat length and nucleotide content for this case’s G_4_C_2_ repeat expansion, but read depth using this approach limited our ability to accurately assess G_4_C_2_ content.

Four of the eight PacBio Sequel reads capturing the *C9orf72* repeat locus were clearly expanded and four were not. Repeat lengths for the four reads capturing the wild-type (non-pathogenic) allele ranged from 46 to 50 nucleotides, where two measured exactly 48 nucleotides (gain of 30 compared to hg38), equating to eight total repeats (Supplemental figure 3c; Figure 6b); these results matched our fragment analysis (Figure 5d). Of the four reads capturing an expanded allele, three did not bridge the entire repeat region, where one captured 178 nucleotides in the repeat region (approximately 30 repeats; Figure. 6c, 7-red; Supplemental data 5), another captured 419 nucleotides (approximately 69 repeats; Figure. 6c, 7-blue; Supplemental data 6), and the third captured 5471 nucleotides (approximately 912 repeats; Figure. 6c, 7-green; Supplemental data 7). It is possible that the read capturing 419 nucleotides bridged the repeat because the end of the read closely matches the sequence adjacent to the repeat region (Supplemental figure 3d). The fourth read, however, spanned the entire repeat with 7941 nucleotides (approximately 1324 repeats; Figure. 6c, 7-brown; Supplemental figure 3e; Supplemental data 8), which falls easily within the Southern blot’s range (Figure 5f). Mean GC content across the case’s repeat region was 87.7% compared to 97.0% for the C9-774 plasmid repeat region. Based on the percentage of plasmid reads that spanned the C9-774 repeat region (43.8%), we would expect approximately half of the reads covering the expanded allele to span at least 774 repeats, which we observed in these data. These data demonstrate the PacBio Sequel is able to sequence through the challenging *C9orf72* G_4_C_2_ repeat, and is well suited for genetic discovery efforts involving large repeat expansions with appropriate sequencing depth.

### Greater read depth through targeted PacBio sequencing enables improved nucleotide content characterization

Because performing long-read, whole-genome sequencing is costly and excessive when investigating a small region, we also tested PacBio’s relatively new No-Amp targeted sequencing method [21] across the *C9orf72* repeat expansion in a case with a smaller repeat (sample 1; Figure 5c). This method allowed us to achieve deeper read depth and more accurately assess nucleotide content compared to the whole-genome approach. We used a sample with a shorter expansion to maximize the number of sequencing passes for individual DNA molecules, thus increasing overall quality of the circular consensus sequences.

We obtained 828 circular consensus sequences for sample 1, where approximately 70% (576 of 828) of reads measured exactly two repeats (the individualvs unexpanded allele; Figure 5a), 14% (115 of 828) were within six nucleotides of two repeats, and 16% (134 of 828) were from expanded alleles. We excluded any sequences that did not read through the entire repeat, determined by alignment (see methods), thus, all included reads represent full-length repeat alleles. The repeat distribution from the expanded alleles suggests mosaicism (by length) with two modes at approximately 110 and 870 repeats (Figure 8). Without prior estimates from the Southern blot (Figure 5c), we likely would have estimated the primary populations of this individual’s repeats at 2,110, and 870 repeats. Because of the Southern blot, however, we know the primary population of the individual’s expanded repeat is near 1000 repeats. The 97.5^th^ and 99^th^ percentiles of the distribution’s probability density function (from the PacBio Sequel) are approximately 964 and 1011 repeats, which closely resemble estimates by Southern blot (Figure 5c).

**Figure 8.**
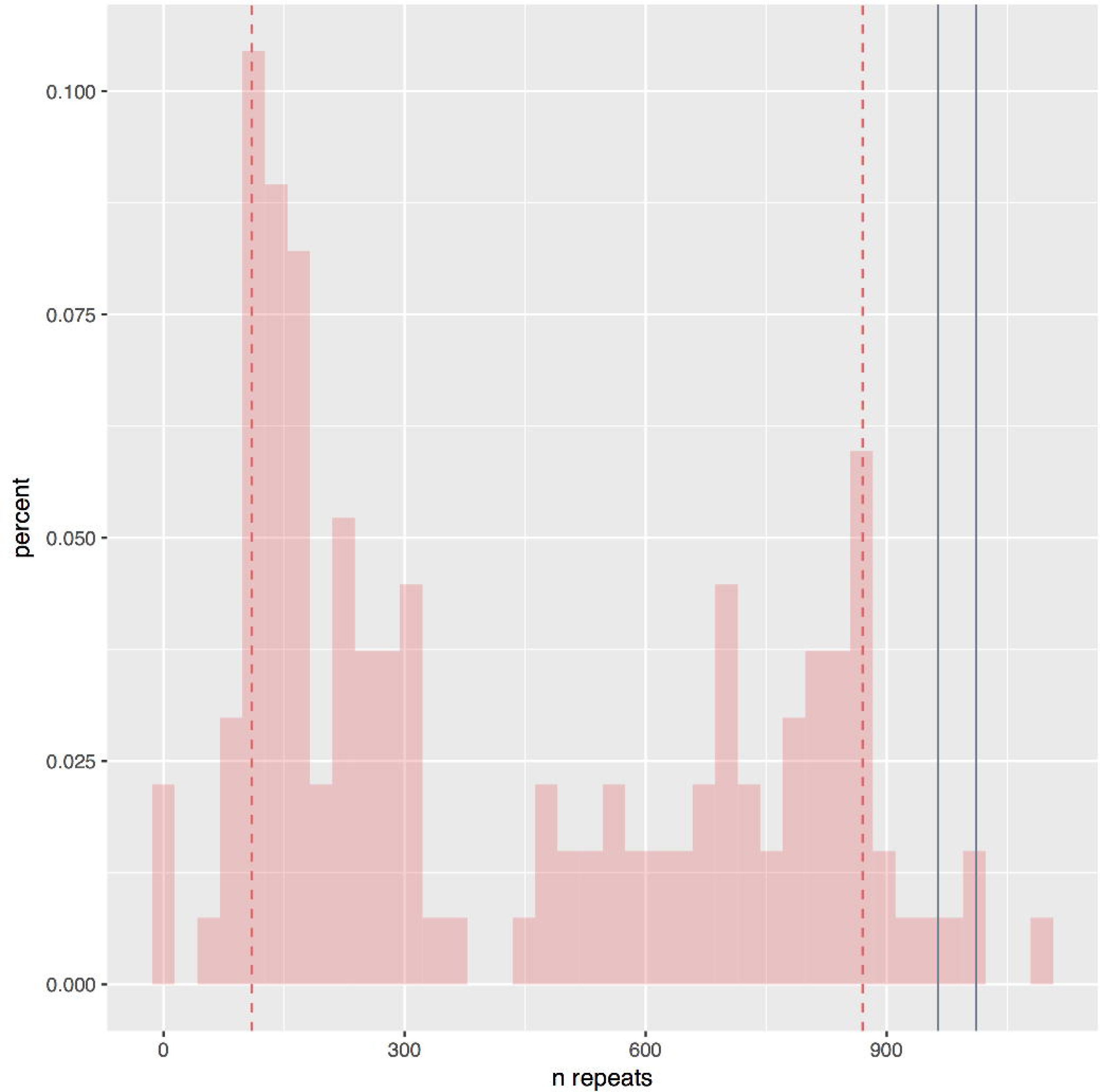
Repeat length distribution using the PacBio no-amplification (No-Amp) targeted sequencing method. We sequenced sample 1 using the PacBio no-amplification (No-Amp) targeted sequencing method and obtained 828 circular consensus sequences. Approximately 70% (576 of 828) of reads covered the individual’s wild-type allele (two repeats), 14% (115 of 828) were within six nucleotides of two repeats, and 16% (134 of 828) were from expanded alleles. The repeat distribution from the expanded alleles shows two modes at approximately 110 and 870 repeats. Without prior estimates from the Southern blot (Figure 5c), we likely would have estimated the primary populations of this individual’s repeats at 2,110, and 870 repeats. Because of the Southern blot, however, we know the primary population of the individual’s expanded repeat is near 1000 repeats, though it is possible the C9orf72 repeat expansion runs artificially high by Southern Blot because of méthylation or high GC content. Assuming the Southern blot is fairly accurate, however, the 97.5th and 99th percentiles of this distribution’s probability density function are approximately 964 and 1011 repeats, which closely resemble estimates by Southern blot (Figure 5c).

Based on the most common simple sequence repeats (SSRs) in reads capturing the repeat expansion, nucleotide content from sample 1 is likely mostly pure G_4_C_2_ repeat. We used PERF (which stands for “PERF is an Exhaustive Repeat Finder”) [25] to measure raw GC content and the most common SSRs found in the repeat region using all expanded circular consensus repeat reads with a minimum read quality of 0.9 and at least two passes around the DNA molecule. Raw GC content within the repeat region was 99.2%. We also found G_4_C_2_ SSR frequency was 81.6%, followed by G_3_C_2_ and G_4_C_1_ with 17.5% and 0.3% frequency, respectively.

Additionally, we performed the Long Amplicon Analysis (LAA2) using SMRTLink 5.1.0 to generate overall consensus sequences across all extended repeat reads and measure the likelihood that any of the non-G_4_C_2_ sequence motifs were real. LAA2 generated one consensus sequence with a predicted accuracy > 99.9% (Supplemental data 9) where the sequence was supported by 301 subreads with an estimated 99.91% accuracy. Looking at only the repeat region, the consensus sequence of the repeat was approximately 894 repeats (5364 nucleotides) with 100% GC and 95.7% G_4_C_2_ content. LAA2 suggested 4.3% of the repeat consisted of G_3_C_2_ interruptions. Some of the other consensus sequences with predicted accuracies > 96% supported small non-GC interruptions. Estimating a consensus sequence in a region with such high mosaicism is not trivial, thus, these results should be interpreted cautiously, but demonstrates the need for a larger study.

Because insertions and deletions (INDELs) are the most common error in PacBio sequencing [26,27], we suspect most or all of the G_3_C_2_ and G_4_C_1_ SSRs are sequencing or basecalling error. Thus, treating all SSR motifs within a Levenshtein distance [28] of one as the expected G_4_C_2_ motif, G_4_C_2_ accounted for 100% of all observed SSRs in the top consensus sequence. Levenshtein distance, which is closely related to the Hamming distance, measures the number of changes between two character sequences, including insertions, deletions, and substitutions. While most of the G_3_C_2_ SSRs are likely false, we cannot rule out that some may be real, potentially affecting repeat-associated non-ATG (RAN) translation [29–31], and disease development and progression. A single G_3_C_2_ interruption would cause a frameshift, resulting in translational transitions from the relatively benign poly(GP) dipeptide repeat to the highly toxic poly(GR) repeat [32–35], or stop a toxic poly(GR), transitioning to the less toxic poly(GA). A larger, deeper sequencing study across the *C9orf72* repeat region in ALS/FTD cases and controls is merited to determine whether there is an association with clinical phenotypes.

## Discussion

Here, we showed that both PacBio and ONT long-read sequencing technologies can sequence through the SCA36 ‘GGCCTG’ and the *C9orf72* G_4_C_2_ repeat expansions in relatively controlled repeats in plasmids, and that the PacBio Sequel can sequence through a human *C9orf72* repeat expansion, in its entirety, depending on length. Additionally, we demonstrated the PacBio No-Amp targeted sequencing method can identify the unexpanded allele and determine whether the individual carries a repeat expansion. These results demonstrate the potential these technologies offer in clinical testing, genetic counseling, and future structural mutation genetic discovery efforts—including those involving challenging repeat expansions. For example, the *C9orf72* G_4_C_2_ repeat expansion could have been discovered years earlier if long-read sequencing technologies had been available. Through great effort, using the best approaches available at the time, the G_4_C_2_ repeat was discovered in 2011 [3,6], approximately five years after chromosome 9p was initially implicated in both ALS and FTD [36,37]. With current long-read sequencing technologies, we can decrease the time to discover and characterize such mutations, begin studying them at the molecular level, and potentially translate them for use in clinical and genetic counseling environments.

We found that both platforms are fully capable of sequencing through challenging repeats like the SCA36 ‘GGCCTG’ and *C9orf72* G_4_C_2_ repeat expansions when cloned into plasmids. It is unclear why the read length distributions for ONT’s MinION were much tighter than those from the PacBio RS II, but it shows promise for future ONT MinION applications. Additionally, while median read lengths were highly similar between the PacBio RS II and ONT MinlON, the MinlON had a higher percentage of reads that extended all the way through the repeat regions for all three repeat plasmids, particularly the C9-423 and C9-774 plasmids.

Both the PacBio RS II and ONT MinION correctly identified the maximal expected number of repeats in the C9-774 plasmid, based on their consensus sequences, but base calling accuracy in the repeat regions was higher for the PacBio RS II. The PacBio RS II attained approximately 99.8% consensus accuracy, while the ONT MinION consensus sequence was only 26.6% accurate because of the mixed nucleotides in the consensus sequence. We believe this will be relatively easy to address in the ONT base calling algorithms because it appears systematic based on the RRRRCM and RRRRMC repeats in the consensus sequences, which demonstrates the same base calling errors occur consistently.

After verifying both PacBio’s and ONT’s technologies were capable of sequencing repeats in plasmids, we tested the technologies’ ability on two *C9orf72* G_4_C_2_ expansion carriers and found the PacBio Sequel is capable of sequencing through challenging GC-rich repeat expansions, but throughput was problematic for the ONT MinION in this study. Newer chemistries and hardware from ONT are likely to alleviate this issue. During the timeline of our study, ONT released the GridlON and PromethlON sequencers that are based on the same nanopore technology and have greater throughput. The PromethION, in particular, can run many more flowcells concurrently, and each individual PromethION flowcell has significantly more nanopores than the MinlON and GridlON flowcells. We anticipate at least the PromethION will be suitable for large repeat studies, based on the MinION’s performance in the plasmids, but we cannot be certain without further testing.

Using the PacBio Sequel, we attained 8x coverage across the *C9orf72* G_4_C_2_ repeat region for sample 2 using the whole-genome approach, where four reads covered the individual’s expected eight-repeat (unexpanded) allele, three reads that ended 30, 69, and 912 repeats into the expansion, respectively, and one read that fully spanned an expanded repeat region with 1324 repeats. The read spanning 1324 repeats is on the lower end of the Southern blot, suggesting longer repeat alleles may have been inaccessible to the PacBio Sequel, perhaps simply because their size impedes loading into the zero-mode waveguide (ZMW) wells. Deeper sequencing is generally required to detect the mutation using a variant caller, but we demonstrate here that the technology is capable of generating such reads, as a proof of principle. We also could not reliably estimate G_4_C_2_ content for sample 2 because of few reads, and each read had only a single sequencing pass. These data do demonstrate, however, that the PacBio Sequel is capable of sequencing through at least a large portion of what may be the most challenging GC-rich repeat expansion known. Discovering whether a structural variant exists, its location, and its general nucleotide makeup is the critical first step to understanding its role in human disease.

Additional studies will be necessary to determine the maximum repeat size that these technologies can span, but our data reiterates the PacBio Sequel is adequate for genetic discovery efforts already [38–41]—and suggests it is capable of sequencing and identifying large repeats. With sufficient read depth, the reads do not necessarily need to bridge the entire repeat (or other large structural variant) to discover whether it exists and characterize the general nucleotide content. Additional experiments can clarify size and nucleotide content, if the sequencing technology was unable to span the variant entirely, or with lower-quality base calls.

After verifying the PacBio Sequel was able to sequence through the *C9orf72* G_4_C_2_ repeat expansion using whole-genome sequencing in a human case, we tested PacBio’s No-Amp targeted sequencing approach to determine how well it can characterize nucleotide content with the increased read depth, and assess how amenable the approach is for clinical and genetic counseling environments. Our results suggest the method can identify an individual’s unexpanded allele, determine whether the individual carries a repeat expansion, and can estimate size up to at least 5kb, though a larger study is needed. While being able to perfectly determine an individual’s expansion size regardless of its length would be ideal, knowing the exact repeat expansion size does not clarify prognosis for *C9orf72* G_4_C_2_ repeat expansion carriers [8], mitigating the need to determine the expansion’s precise size. Additionally, the repeat size is known to be highly variable throughout various body tissues, including different brain regions, and even within a small tissue piece from the same brain region [3,8,42–44]. This is further demonstrated by the smear within the Southern blots for both symptomatic carriers included in this study.

While repeat size is not informative for prognosis, being able to assess overall G_4_C_2_ content and detect repeat interruptions may be informative for prognosis, but more information is required. We were able to more accurately assess G_4_C_2_ content using the targeted approach, though distinguishing between G_4_C_2_ and G_3_C_2_ motifs is likely unreliable at this stage. Treating all G_3_C_2_ motifs as G_4_C_2_, we estimate the G_4_C_2_ content for sample 1 is > 99%, and potentially 100%. There is some evidence supporting potential GaC_2_ and non-GC interruptions, but experimentally verifying these finer differences in low-complexity repeat regions is nontrivial. A larger study will be important to determine whether it is possible to identify more pronounced interruptions, or even distinguishing between G_4_C_2_ and G_3_C_2_. The ability to identify an individual’s unexpanded allele, clearly indicate expansion status, and assess nucleotide content in a single experiment could have important implications in clinical and genetic counseling environments, and will certainly be investigated further in the research environment.

Existing challenges for the No-Amp targeted sequencing method include low throughput, and it inherently selects for shorter reads, or repeats in this case, because of both loading bias (shorter fragments load preferentially) and that the polymerase is less likely to traverse longer repeats as reliably as shorter repeats. This is likely why there is a statistical mode at approximately 110 repeats (Figure 8), even though there is no observable band at that size in the Southern blot. We are confident the reads are real, however, as the Southern blot clearly shows size mosaicism, and the adjacent sequence on both sides of the repeat region aligned on both sides of the repeat region for each read with ≥85% identity for all included reads. The bias towards shorter reads does misrepresent the primary size distribution, but we were still able to determine that the individual carries a repeat expansion, and we were able to accurately estimate the size in this case. Determining whether an individual carries a repeat expansion in an automated fashion would be relatively simple using this approach.

Given that several studies have shown the repeat expansion is variable across tissues within a given patient [8,42–44], we suggest that a large, deep long-read sequencing study across the *C9orf72* repeat is important to better understand how repeat content affects disease onset and progression. Repeat interruptions are known to mitigate disease in other neurodegenerative disorders [14–17]. Fully characterizing the repeat at the nucleotide level in a large cohort may have critical implications on our understanding of disease etiology, development, duration, and on future therapy. A large, long-read sequencing study of affected *C9orf72* G_4_C_2_ repeat expansion carriers would also allow us to characterize mosaicism within individuals; there may be expansion sub-species that explain the more aggressive forms of ALS and FTD, that are not measurable through traditional methods, such as Southern blotting.

Cost is a limiting factor for long-read sequencing technologies, often making it impractical for large studies or for diagnostic use. Because of these limitations, researchers have made great efforts to maximize the utility of short-read sequencing technologies, employing the large amount of short-read sequencing data already generated across nearly every disease currently studied. An excellent example is the effort to identify repeat expansions based on evidence in existing short-read data [45,46]. These efforts offer researchers that have already generated short-read data for individuals the ability to determine whether an individual has a repeat expansion, but only if the repeat expansion and its location are already known. The approaches are also generally unable to estimate the repeat size. The limitations in these approaches reflect the limitations of short-read sequencing, because short reads cannot span even relatively small repeat expansions. Long-read sequencing, while having a much higher error rate, addresses these limitations, and may be more amenable to regular use in the future. For now, long-read sequencing may be ideal for small familial studies or for smaller studies intent on identifying repeat expansions that exist among a small cohort of cases. Researchers could then follow up with more cost-effective methods such as repeat-primed PCR or Southern blotting.

Knowing PacBio and ONT long-read sequencing technologies are fully capable of sequencing through challenging disease-causing repeats, such as the SCA36 ‘GGCCTG’ and *C9orf72* ‘GGGGCC’ repeats, lays important ground work for future sequencing studies to understand the nucleotide-level nature of all repeat-expansion disorders. It also demonstrates that long-read sequencing technologies offer great potential for future repeat expansion discovery efforts, and may be useful in clinical and genetic counseling environments for either the *C9orf72* repeat expansion specifically, or for other large structural mutations; the ability to target specific regions will be particularly important in several settings. Further utilizing these technologies in larger studies will be critical to properly characterizing known repeats (e.g., *C9orf72)* and their allelic distributions (size and content) on the nucleotide level to better understand how they contribute to disease.

## Conclusions

Our results demonstrate that long-read sequencing is well suited to characterizing known repeat expansions, and for discovering new disease-causing, disease-modifying, or risk-modifying repeat expansions that have gone undetected with conventional short-read sequencing. These results have important implications on future genetic discovery efforts, as many diseases are caused by repeat expansions or other large structural variants. Larger studies focusing on the *C9orf72* expansion in ALS and FTD will be important to determine heterogeneity and whether the repeats are interrupted by non-G_4_C_2_ content. Such interruptions are likely to modify the disease course or age of onset, as shown in other repeat-expansion disorders [14–17]. These results have broad implications across all diseases where the genetic etiology remains unclear.

## Materials and Methods

### Study participants

Cerebellar samples included in this study were obtained from the Mayo Clinic Brain Bank, following Mayo Clinic’s IRB protocols. Sample 1 was female and was diagnosed with FTD with an age of onset and duration of approximately 44 and 12 years, respectively. Sample 2 was male, and was diagnosed with FTD with an age of onset and duration of 70 and 2 years, respectively. Fluorescent and repeat-primed PCR [3], and Southern blot [24] techniques were previously described.

### Repeat plasmids

To generate the G_4_C_2_ repeat 423 and 774 expression vectors, we used muscle or spleen DNA from an affected *C9orf72* expansion carrier as a template in a nested PCR strategy. We used ThermalAce DNA Polymerase (Invitrogen) to amplify the 66 G_4_C_2_ repeat region of a previously constructed G_4_C_2_ repeat plasmid, including 113 and 99 nucleotides of 5’ and 3’ flanking sequence, respectively. We then used these intermediate plasmids to construct PCR products for the 423 and 774 sequence that were subsequently cloned into the pAG3 and pcDNA6 expression vectors containing 3 different C-terminal tags in alternate frames. The *EGFP* gene and SCA36 ‘GGCCTG’ 66 repeat were cloned into pAAV expression vectors. The SCA36 clone was Sanger sequenced to determine the number of repeats, but the others were too long for Sanger sequencing.

### Sequencing

Plasmid sequencing libraries were prepared by first linearizing plasmids individually and then pooling in equal concentration based on NanoDrop (ThermoFisher Scientific) measurements. Plasmids were linearized using the restriction site that provided the most non-repeat sequence up and downstream of the repeat, itself. EGFP, SCA36, and C9-774 plasmids were linearized using the Avrll restriction enzyme, while C9-423 was linearized using Mlul (Figure 1). DNA was then purified using Agencourt AMPure XP beads, per the manufacturer’s recommended protocol, and split equally for ONT MinION and PacBio RS II library preparation.

We used the ONT MinlON “ID Genomic DNA by ligation” kit and protocol (SQK-LSK108) for MinION library preparation and sequencing, skipping optional DNA fragmentation and DNA repair steps to maximize DNA size and quantity, respectively. Briefly, we began with end repair and dA-tailing using all recommended reagents and steps, which include mixing the DNA, Ultra II end-prep reaction buffer, Ultra II end-prep enzyme mix, and nuclease-free water, and incubating at 20°C for five minutes and 65°C for five minutes. DNA was then purified using Agencourt AMPure XP beads, and eluted in nuclease-free water. The 1D sequencing adapter was then ligated by mixing the DNA, 1D adapter mix (AMX1D), and Blunt/TA ligation master mix and incubating for 10 minutes. DNA was then purified again using the Agencourt AMPure XP beads, but washing with the ONT Adapter Bead Binding (ABB) buffer and eluting in ONT’s elution buffer (ELB). Flow cells were primed and loaded per recommended procedure and sequenced using the 48-hour sequencing protocol in MinKNOW. We performed the recommended quality control run on all flow cells prior to priming and loading the library and only used flow cells that had >1000 active pores and did not have existing air bubbles in the Application-Specific Integrated Circuit (ASIC) upon opening.

Plasmid libraries for the PacBio RS II were generated following PacBio’s protocol for the SMRTbell Template Prep Kit 1.0 (Part #100-259-100) and PacBio’s “Procedure & Checklist—10 kb Template Preparation and Sequencing”. Briefly, DNA damage repair, end repair, blunt end hairpin adapter ligation, and final exonuclease treatment were performed using 5μg of intact, non-sheared pooled plasmid DNA. AMPure PB magnetic beads (Pacific Biosciences) were used for all purification steps. Qualitative and quantitative analysis were performed using Advanced Analytical Fragment Analyzer (AATI) and Qubit fluorometer with Quant-iT dsDNA BR Assay Kits (Invitrogen). SMRTbell templates were annealed to v2 sequencing primers then bound to DNA polymerase P6 following PacBio’s protocol using the DNA/Polymerase Binding Kit P6 (part #: 100-356-300), as directed using Binding Calculator version 2.3.1.1. Polymerase-template complexes were purified per manufacturer’s protocol (PacBio) using Pacific Biosciences Magbead Binding Kit (part #: 100-133-600) and setting up sample reaction as directed using Binding Calculator. Sequencing was carried out on the PacBio RS II (SMRT) sequencer, equipped with MagBead Station upgrade, using C4 DNA Sequencing Kit 2.0 (Part #: 100-216-400) reagents. The sample was loaded onto a single SMRT cell v3 (part #: 100-171-800) and the movie length was 360 min. Secondary Analysis was performed using Pacific Biosciences SMRT Portal, using SMRT Analysis System software (v2.3.0) for Sub-read filtering.

When sequencing the affected *C9orf72* repeat expansion carrier, we extracted DNA from the affected *C9orf72* repeat expansion carrier using the Agilent RecoverEase DNA Isolation Kit (Agilent Technologies) and a standard, previously-described isolation protocol [8]. We followed the same ONT MinlON “ID Genomic DNA by ligation” kit and protocol used for the plasmids, using 15 total flow cells. For PacBio whole-genome sequencing, we sequenced the DNA using the Sequel instead of the RS II because the Sequel provides greater throughput. We prepared the PacBio Sequel library per PacBio’s recommended protocol “Procedure & Checklist—20kb Template Preparation Using BluePippin Size-Selection System”, using the Megaruptor 2 (Diagenode, Denville, NJ, USA) for shearing and the Fragment Analyzer (Advanced Analytical, Ankeny, IA, USA) to size the DNA. To summarize, DNA was sheared to 35kb on the Megaruptor 2 and prepared using the SMRTbell Template Prep Kit 1.0-SPv3 (part #: 100-991-900). Qualitative and quantitative analysis were performed using Advanced Analytical Fragment Analyzer (AATI) and Qubit fluorometer with Quant-iT dsDNA BR Assay Kits (Invitrogen). SMRTbell templates were annealed to v3 sequencing primers then bound to DNA polymerase 2.0 following PacBio’s protocol using the Sequel Binding and Internal Ctrl Kit 2.0 (part #: 101-400-900). Excess polymerase was removed from the binding reaction using the PacBio Loading Cleanup Bead Kit (part #: 100-715-300). MagBead binding was performed using the PacBio protocol for kit 100-125-900. Cleaned samples were loaded onto five SMRTcells with Sequel Sequencing Kit 2.0 following PacBio recommendations, and sequenced on the PacBio Sequel (SMRT) sequencer.

The PacBio No-Amp targeted sequencing procedure, currently in development, uses the CRISPR-Cas9 system to target and enrich a region of interest without PCR amplification (Figure 9) [21]. For sample 1, 20μg non-sheared, genomic DNA was digested with high fidelity restriction enzyme EcoRI-HF (New England Biolabs, PN R3101S) to excise the target region. A SMRTbell library was prepared from the EcoRI-HF digested products by ligation with a hairpin adapter containing an overhang sequence complementary to the EcoRI-HF cut site using *E. coli* DNA ligase (New England Biolabs, PN M0205S). Genome complexity reduction was then performed by incubating each sample with high fidelity restriction enzymes KpnI-HF and SpeI-HF (New England Biolabs, PN R3142S and R3133S, respectively) and Exonuclease III and VII (Pacific Biosciences, part of SMRTbell Template Prep Kit 1.0, PN 100-259-100). Up to 1 μg of the complexity-reduced SMRTbell library was subjected to Cas9 digestion with a single guide RNA specific to sequence adjacent to the target region. Oligos comprising the guide RNA (crRNA and tracrRNA) were obtained from Integrated DNA Technologies containing an Alt-R modification to prevent RNase degradation. Cas9 Nuclease, *S. pyogenes*, was obtained from New England Biolabs (PN M0386S). Cas9-digested SMRTbell templates were ligated with a poly(A) hairpin adapter using T4 DNA ligase (Pacific Biosciences, part of SMRTbell Template Prep Kit 1.0, PN 100-259-100) producing asymmetric SMRTbell templates. Failed ligation products were removed by treatment with Exonuclease III and VII (Pacific Biosciences). Asymmetric SMRTbell templates were then enriched with MagBeads and buffers from MagBead Kit v2 (Pacific Biosciences, PN 100-676-500). In preparation for sequencing, a standard PacBio sequencing primer, lacking a poly(A) sequence, was annealed to the enriched SMRTbell templates in diluted Primer Buffer v2 (Pacific Biosciences, PN 001-560-849). Sequel DNA Polymerase 2.1 was bound to the primer-annealed SMRTbell templates with reagents from the associated Sequel Binding Kit 2.1 (Pacific Biosciences, PN 101-365-900). The sample complex was purified with a modified AMPure PB purification protocol (Pacific Biosciences, PN 100-265-900). The entire purified sample complex was loaded on a single SMRT Cell (Pacific Biosciences, PN 101-008-000) and sequenced on a Sequel System using an immobilization time of 4 hours and movie time of 10 hours.

**Figure 9.**
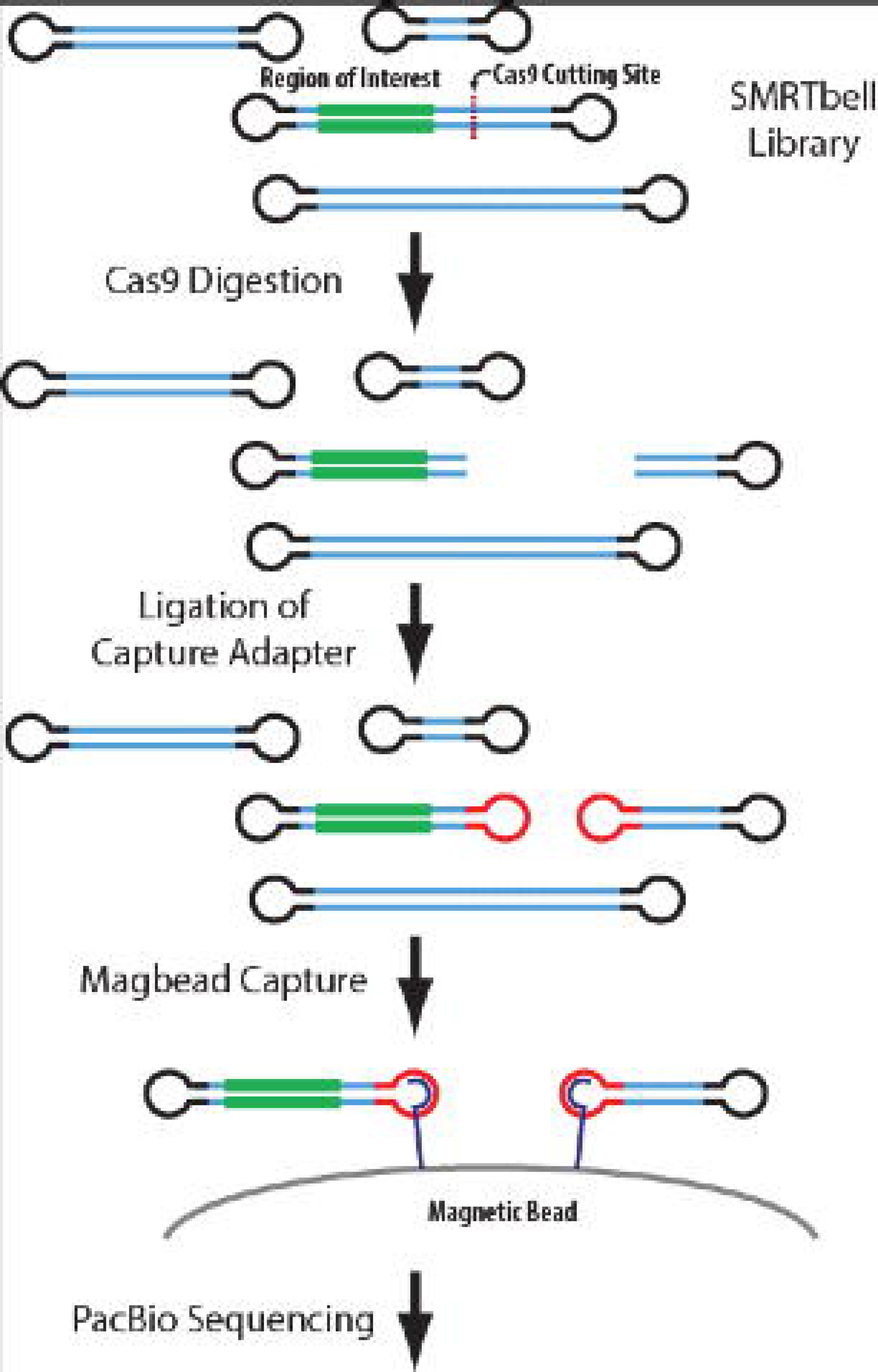
Schematic of PacBio no-amplification (No-Amp) targeted sequencing. We applied the PacBio no-amplification (No-Amp) Targeted Sequencing method to a *C9orf72* G_4_C_2_ repeat expansion carrier to better characterize the repeat’s nucleotide content. The No-Amp targeted sequencing method begins with typical SMRTbell library preparation after the target region is excised by restriction enzyme digestion. Cas9 digestion follows with a guide RNA specific to sequence adjacent (Cas9 Cutting Site) to the region of interest (green), leaving the SMRTbells blunt ended. In this case, the guide RNA was specific to sequence upstream (5’) of the G_4_C_2_ repeat expansion on the anti-sense strand. A new capture adapter (red) is then ligated to the blunt ends and captured using magnetic beads (magbeads). This process enriches the library for reads containing the region of interest to maximize read depth.

### Analyses

#### Base calling and alignment

All ONT MinION reads were base called using Albacore version 1.2.1 and PacBio reads were base called using SMRT Analysis version 2.3 for the PacBio RS II data and SMRT Link version 4.0 for the PacBio Sequel data. After base calling, plasmid reads were separated by sequence unique to each plasmid backbone (Supplemental data 10) using BLAST version 2.6.0. Reads were first separated by backbone to avoid mixing reads from plasmids with the same repeat (e.g., C9-423 and C9-774 had the same repeat, but different backbones). EGFP and SCA36 were then separated by their unique sequences, the *EGFP* gene and the ‘GGCCTG’ repeat, respectively. Separated reads were then aligned to their respective reference sequences using graphmap version 0.5.1 [47] with default parameters. All whole-genome reads from sample 1 were aligned to human reference genome hg38 using graphmap version 0.5.1 with default parameters. Reads across the *C9orf72* repeat expansion locus for the affected expansion carrier were hand curated, using the original graphmap alignment as a guide, where specific sequence “landmarks” were used to define the repeat region. The final hand-curated alignments are included in Supplemental figure 3. Alignments were visualized using the Integrative Genomics Viewer (IGV) version 2.3.94 [48,49].

Circular consensus (CCS) reads from the PacBio No-Amp targeted sequencing method were generated using the “ccs” binary in SMRTLink version 5.1.0, using all subreads with 0 or more passes and predicted accuracy ≥ 80%. On-target reads were identified by aligning the adjacent 100 nucleotides from both sides of the *C9orf72* repeat sequence in hg38 (Supplemental data 11) using the Smith-Waterman [50] local alignment algorithm. The repeat region was defined as the nucleotides between the alignments of both adjacent sequences, where we required ≥ 85% similarity. We excluded any sequences that did not read through the repeat. We used the “laa” binary in SMRTLink version 5.1.0 to generate overall consensus sequences across all on-target subreads. We used PERF [25] to estimate GC content and simple sequence repeat (SSR) frequencies, except we calculated the frequencies as the total number of times a given SSR motif (e.g., G_4_C_2_) was found in the sequence (without overlap); PERF counts a set of repeats (e.g., 100 consecutive, uninterrupted G_4_C_2_ motifs) as a single occurrence.

#### Statistical analyses

Total read counts, reads extending through the repeat, and repeat lengths for each plasmid were calculated using pysam version 0.12, which is a wrapper for samtools [51]. To calculate reads extending through the repeats, we required the read to have sequence within 100 nucleotides before and after the repeat region, based on its alignment (i.e., the read could not start or end within the repeat). Repeat lengths were calculated by extracting all bases that aligned within the reference sequence’s repeat region, removing all gaps, and counting remaining nucleotides. Summary statistics, including median read lengths, and plots were generated in R version 3.4.1 [52] and ggplot2 version 2.2.1 [53]. Coverage across the *C9orf72* repeat expansion carrier’s genome was calculated using bedtools genomecov [54].

To calculate error rates for ONT MinION and PacBio RS II in the C9-774 plasmid repeat region, we extracted individual repeat sequence from each read, as previously explained, and aligned them to the reference repeat sequence equal to the measured repeat length of the given read. We used the global Needleman-Wunsch algorithm [23] for these alignments, for the most accurate result.

**Abbreviations**
ALS: Amyotrophic Lateral Sclerosis
FTD: Frontotemporal Dementia
G_4_C_2_: GGGGCC
PacBio: Pacific Biosciences
ONT: Oxford Nanopore Technologies
SCA36: spinocerebellar ataxia type 36

## Declarations

### Ethical Approval and Consent to participate

The Mayo Clinic Institutional Review Board (IRB) approved all procedures for this study and we followed all appropriate protocols.

### Consent for publication

All participants were properly consented for this study.

### Availability of supporting data

All data are available upon reasonable request to the corresponding author.

## Authors’ contributions

ME, LP, and JF developed and designed the study, and wrote the manuscript. ME, BB, and MS performed analyses. SF, JS, KJW, TG, MP, IM, MDH, PB, and MvB performed necessary experiments. MvB and RR provided expansion status on samples. DD provided tissue from the Mayo Clinic Brain Bank and the pathology report. All authors read and approved the final manuscript.

## Acknowledgements

We gratefully acknowledge contributions by the Brigham Young University DNA Sequencing Center (DNASC), and the Mayo Clinic Molecular Biology Core who provided valuable input and services. We also recognize the Mayo Clinic Research Computing Services who provided the requisite computational resources for all analyses performed in this study.

## References

1. La Spada AR, Taylor JP. Repeat expansion disease: progress and puzzles in disease pathogenesis. Nat Rev Genet. 2010;11:247–58.

2. Orr HT, Chung M, Banfi S, Kwiatkowski TJ, Servadio A, Beaudet AL, et al. Expansion of an unstable trinucleotide CAG repeat in spinocerebellar ataxia type 1. Nat Genet. 1993;4:221–6.

3. DeJesus-Hernandez M, Mackenzie IR, Boeve BF, Boxer AL, Baker M, Rutherford NJ, et al. Expanded GGGGCC hexanucleotide repeat in non-coding region of C9ORF72 causes chromosome 9p-linked frontotemporal dementia and amyotrophic lateral sclerosis. Neuron. 2011;72:245–56.

4. Kieleczawa J. Fundamentals of Sequencing of Difficult Templates—An Overview. J Biomol Tech JBT. 2006;17:207–17.

5. Zhao X, Haqqi T, Yadav SP. Sequencing telomeric DNA template with short tandem repeats using dye terminator cycle sequencing. J Biomol Tech JBT. 2000;11:111–21.

6. Renton AE, Majounie E, Waite A, Simón-Sánchez J, Rollinson S, Gibbs JR, et al. A Hexanucleotide Repeat Expansion in C9ORF72 Is the Cause of Chromosome 9p21-Linked ALS-FTD. Neuron. 2011;72:257–68.

7. van Blitterswijk M, DeJesus-Hernandez M, Rademakers R. How do C90RF72 repeat expansions cause ALS and FTD: can we learn from other non-coding repeat expansion disorders? Curr Opin Neurol. 2012;25:689–700.

8. van Blitterswijk M, DeJesus-Hernandez M, Niemantsverdriet E, Murray ME, Heckman MG, Diehl NN, et al. Association between repeat sizes and clinical and pathological characteristics in carriers of C9ORF72 repeat expansions (Xpansize-72): a cross-sectional cohort study. Lancet Neurol. 2013;12:978–88.

9. Van Mossevelde S, van der Zee J, Cruts M, Van Broeckhoven C. Relationship between C9orf72 repeat size and clinical phenotype. Curr Opin Genet Dev. 2017;44:117–24.

10. Nakano M, Okumura N, Nakagawa H, Koizumi N, Ikeda Y, Ueno M, et al. Trinucleotide Repeat Expansion in the TCF4 Gene in Fuchs’ Endothelial Corneal Dystrophy in Japanese. Invest Ophthalmol Vis Sci. 2015;56:4865–9.

11. Mahadevan M, Tsilfidis C, Sabourin L, Shutler G, Amemiya C, Jansen G, et al. Myotonic Dystrophy Mutation: An Unstable CTG Repeat in the 3⟀ Untranslated Region of the Gene. Science. 1992;255:1253–5.

12. Campuzano V, Montermini L, Moltà MD, Pianese L, Cossée M, Cavalcanti F, et al. Friedreich’s ataxia: autosomal recessive disease caused by an intronic GAA triplet repeat expansion. Science. 1996;271:1423–7.

13. Verkerk AJMH, Pieretti M, Sutcliffe JS, Fu Y-H, Kuhl DPA, Pizzuti A, et al. Identification of a gene (FMR-1) containing a CGG repeat coincident with a breakpoint cluster region exhibiting length variation in fragile X syndrome. Cell. 1991;65:905–14.

14. Kraus-Perrotta C, Lagalwar S. Expansion, mosaicism and interruption: mechanisms of the CAG repeat mutation in spinocerebellar ataxia type 1. Cerebellum Ataxias [Internet]. 2016; 3. Available from: http://www.ncbi.nlm.nih.gov/pmc/articles/PMC5118900/

15. Matsuura T, Fang P, Pearson CE, Jayakar P, Ashizawa T, Roa BB, et al. Interruptions in the Expanded ATTCT Repeat of Spinocerebellar Ataxia Type 10: Repeat Purity as a Disease Modifier? Am J Hum Genet. 2006;78:125–9.

16. Sakamoto N, Larson JE, Iyer RR, Montermini L, Pandolfo M, Wells RD. GGA*TCC-interrupted triplets in long GAA*TTC repeats inhibit the formation of triplex and sticky DNA structures, alleviate transcription inhibition, and reduce genetic instabilities. J Biol Chem. 2001;276:27178–87.

17. Stolle CA, Frackelton EC, McCallum J, Farmer JM, Tsou A, Wilson RB, et al. Novel, complex interruptions of the GAA repeat in small, expanded alleles of two affected siblings with late-onset Friedreich ataxia. Mov Disord Off J Mov Disord Soc. 2008;23:1303–6.

18. Fratta P, Mizielinska S, Nicoll AJ, Zloh M, Fisher EMC, Parkinson G, et al. *C9orf72* hexanucleotide repeat associated with amyotrophic lateral sclerosis and frontotemporal dementia forms RNA G-quadruplexes. Sci Rep. 2012;2:srep01016.

19. Haeusler AR, Donnelly CJ, Periz G, Simko EAJ, Shaw PG, Kim M-S, et al. C9orf72 nucleotide repeat structures initiate molecular cascades of disease. Nature. 2014;507:195–200.

20. Kobayashi H, Abe K, Matsuura T, Ikeda Y, Hitomi T, Akechi Y, et al. Expansion of intronic GGCCTG hexanucleotide repeat in NOP56 causes SCA36, a type of spinocerebellar ataxia accompanied by motor neuron involvement. Am J Hum Genet. 2011;89:121–30.

21. Tsai Y-C, Greenberg D, Powell J, Hoijer I, Ameur A, Strahl M, et al. Amplification-free, CRISPR-Cas9 Targeted Enrichment and SMRT Sequencing of Repeat-Expansion Disease Causative Genomic Regions. bioRxiv. 2017;203919.

22. No-Amp Targeted Sequencing [Internet]. PacBio. [cited 2018 May 30]. Available from: https://www.pacb.com/applications/targeted-sequencing/no-amp-targeted-sequencing/

23. Needleman SB, Wunsch CD. A general method applicable to the search for similarities in the amino acid sequence of two proteins. J Mol Biol. 1970;48:443–53.

24. van Blitterswijk M, Baker MC, DeJesus-Hernandez M, Ghidoni R, Benussi L, Finger E, et al. C90RF72 repeat expansions in cases with previously identified pathogenic mutations. Neurology. 2013;81:1332–41.

25. Avvaru AK, Sowpati DT, Mishra RK. PERF: An Exhaustive Algorithm for Ultra-Fast and Efficient Identification of Microsatellites from Large DNA Sequences. Bioinformatics [Internet], [cited 2017 Nov 22]; Available from: https://academic.oup.com/bioinformatics/advance-article/doi/10.1093/bioinformatics/btx721/4600186

26. Ross MG, Russ C, Costello M, Hollinger A, Lennon NJ, Hegarty R, et al. Characterizing and measuring bias in sequence data. Genome Biol. 2013;14:R51.

27. Laehnemann D, Borkhardt A, McHardy AC. Denoising DNA deep sequencing data—high-throughput sequencing errors and their correction. Brief Bioinform. 2016;17:154–79.

28. Levenshtein VI. Binary Codes Capable of Correcting Deletions, Insertions and Reversals. Sov Phys Dokl. 1966;10:707.

29. Zu T, Gibbens B, Doty NS, Gomes-Pereira M, Huguet A, Stone MD, et al. Non-ATG-initiated translation directed by microsatellite expansions. Proc Natl Acad Sci USA. 2011;108:260–5.

30. Cleary JD, Ranum LPW. Repeat-associated non-ATG (RAN) translation in neurological disease. Hum Mol Genet. 2013;22:R45–51.

31. Green KM, Linsalata AE, Todd PK. RAN translation-What makes it run? Brain Res. 2016;1647:30–42.

32. Kwon I, Xiang S, Kato M, Wu L, Theodoropoulos P, Wang T, et al. Poly-dipeptides encoded by the C9orf72 repeats bind nucleoli, impede RNA biogenesis, and kill cells. Science. 2014;345:1139–45.

33. Jovičić A, Mertens J, Boeynaems S, Bogaert E, Chai N, Yamada SB, et al. Modifiers of C9orf72 dipeptide repeat toxicity connect nucleocytoplasmic transport defects to FTD/ALS. Nat Neurosci. 2015;18:1226–9.

34. Tao Z, Wang H, Xia Q, Li K, Li K, Jiang X, et al. Nucleolar stress and impaired stress granule formation contribute to C9orf72 RAN translation-induced cytotoxicity. Hum Mol Genet. 2015;24:2426–41.

35. Zhang Y-J, Gendron TF, Ebbert MTW, O’Raw AD, Yue M, Jansen-West K, et al. Poly(GR) impairs protein translation and stress granule dynamics in C9orf72 -associated frontotemporal dementia and amyotrophic lateral sclerosis. Nat Med. 2018;1.

36. Morita M, Al-Chalabi A, Andersen PM, Hosler B, Sapp P, Englund E, et al. A locus on chromosome 9p confers susceptibility to ALS and frontotemporal dementia. Neurology. 2006;66:839–44.

37. Vance C, Al-Chalabi A, Ruddy D, Smith BN, Hu X, Sreedharan J, et al. Familial amyotrophic lateral sclerosis with frontotemporal dementia is linked to a locus on chromosome 9p13.2-21.3. Brain J Neurol. 2006;129:868–76.

38. Aneichyk T, Hendriks WT, Yadav R, Shin D, Gao D, Vaine CA, et al. Dissecting the Causal Mechanism of X-Linked Dystonia-Parkinsonism by Integrating Genome and Transcriptome Assembly. Cell. 2018;172:897–909.e21.

39. Cartwright JF, Anderson K, Longworth J, Lobb P, James DC. Highly sensitive detection of mutations in CHO cell recombinant DNA using multi-parallel single molecule real-time DNA sequencing. Biotechnol Bioeng. 2018;115:1485–98.

40. Lodé L, Ameur A, Coste T, Ménard A, Richebourg S, Gaillard JB, et al. Single-molecule DNA sequencing of acute myeloid leukemia and myelodysplastic syndromes with multiple TP53 alterations. Haematologica. 2017;haematol.2017.176719.

41. Turner TR, Hayhurst JD, Hayward DR, Bultitude WP, Barker DJ, Robinson J, et al. Single molecule real-time DNA sequencing of HLA genes at ultra-high resolution from 126 International HLA and Immunogenetics Workshop cell lines. HLA. 2018;91:88–101.

42. Suh E, Lee EB, Neal D, Wood EM, Toledo JB, Rennert L, et al. Semi-automated quantification of C9orf72 expansion size reveals inverse correlation between hexanucleotide repeat number and disease duration in frontotemporal degeneration. Acta Neuropathol (Berl). 2015;130:363–72.

43. Dols-Icardo O, García-Redondo A, Rojas-García R, Sánchez-Valle R, Noguera A, Gómez-Tortosa E, et al. Characterization of the repeat expansion size in C9orf72 in amyotrophic lateral sclerosis and frontotemporal dementia. Hum Mol Genet. 2014;23:749–54.

44. Nordin A, Akimoto C, Wuolikainen A, Alstermark H, Jonsson P, Birve A, et al. Extensive size variability of the GGGGCC expansion in C9orf72 in both neuronal and non-neuronal tissues in 18 patients with ALS or FTD. Hum Mol Genet. 2015;24:3133–42.

45. Dolzhenko E, Vugt JJFA van, Shaw RJ, Bekritsky MA, Blitterswijk M van, Narzisi G, et al. Detection of long repeat expansions from PCR-free whole-genome sequence data. Genome Res. 2017;gr.225672.117.

46. Dashnow H, Lek M, Phipson B, Halman A, Davis M, Lamont P, et al. STRetch: detecting and discovering pathogenic short tandem repeats expansions. bioRxiv. 2017;159228.

47. Sović I, Šikić M, Wilm A, Fenlon SN, Chen S, Nagarajan N. Fast and sensitive mapping of nanopore sequencing reads with GraphMap. Nat Commun. 2016;7:ncommsll307.

48. Robinson JT, Thorvaldsdóttir H, Winckler W, Guttman M, Lander ES, Getz G, et al. Integrative genomics viewer [Internet]. Nat. Biotechnol. 2011 [cited 2018 May 23]. Available from: https://www.nature.com/articles/nbt.1754

49. Thorvaldsdóttir H, Robinson JT, Mesirov JP. Integrative Genomics Viewer (IGV): high-performance genomics data visualization and exploration. Brief Bioinform. 2013;14:178–92.

50. Smith TF, Waterman MS. Identification of common molecular subsequences. J Mol Biol. 1981;147:195–7.

51. Li H, Handsaker B, Wysoker A, Fennell T, Ruan J, Homer N, et al. The Sequence Alignment/Map format and SAMtools. Bioinforma Oxf Engl. 2009;25:2078–9.

52. R Development Core Team. R: A Language and Environment for Statistical Computing [Internet]. Vienna, Austria: R Foundation for Statistical Computing; 2011. Available from: http://www.R-project.org/

53. ggplot2 - Elegant Graphics for Data Analysis | Hadley Wickham | Springer [Internet], [cited 2017 Sep 15]. Available from: http://www.springer.com/us/book/9780387981413

54. Quinlan AR, Hall IM. BEDTools: a flexible suite of utilities for comparing genomic features. Bioinformatics. 2010;26:841–2.

